# Inhibition of CaOx crystals by *Neolamarckia cadamba*: An *in vivo* approach

**DOI:** 10.1101/253179

**Authors:** P.V Prathibhakumari, G. Prasad

## Abstract

The objective of the study was to find out the effectiveness of the methanol fruit extract of *N. cadamba* on calcium oxalate induced nephrolithiasis in wistar albino rats. Animals were divided into nine groups (n=4) in which group 1 as control, group II as antilithiatic control and group III as lithiatic control. Dose for the methanol fruit extract was selected for the study as 200 and 400mg/kg body weight of fruit extract. Group IV and V were considered as post treatment groups and group VI to IX were co treatment groups. Ethylene glycol in drinking water was given to group II - IX for the induction of renal calculi. All the stone forming constituents such as urea, uric acid and creatinine were reduced significantly (p<0.01) in the extract treated groups. Calcium, oxalate and phosphorous concentrations in kidney were found to be diminished by the supplementation of extract. ICP-MS analysis, Histopathology, microcrystal study and pizzalato’s staining confirmed the efficacy of the fruit extract. In conclusion, the results suggested that the fruit extract is endowed with the property of an antilithiatic drug.

## Introduction

Nephrolithiasis is a complex physicochemical event which starts with urinary supersaturation and then leads to nucleation, growth, aggregation and retention of crystal forming constituents within the kidneys (Butterweck and Khan, 2009) and is a significant health problem in all societies. Several epidemiological data and clinical studies have reported that calcium oxalate (CaOx) followed by calcium phosphate are the frequently encountered type of stones in majority of kidney stones (Daudon *et al*., 1993). Nephrolithiasis is still considered a mysterious disease even though extensive research has been carried out in urology. Various therapies, treatments, investigations and sophisticated instruments are failed to trace out the exact cause and mechanisms of urolithiasis (Pareta *et al*., 2011). Several drugs have been used against nephrolithiasis in solitary or in combination with other drugs. In spite of numerous developments in the treatment of urolithiasis there is no satisfactory drug in modern medicine to remove the stones. Therefore, research gives much attention for better medical therapy and to develop a satisfactory drug to prevent stone formation in lithiatic patients. The understanding of the pathophysiology of renal lithiasis has great importance in the development of effective and safe anti-nephrolithiatic agents.

*Neolamarckia cadamba* is a fast growing tree belongs to the family Rubiaceae (Divyakant *et al*., 2011). The fruits are small round balls, hard and yellow when ripe with sweet taste and are edible. *N. cadamba* is used for the blood, cough, uterine complaints, urinary ailments, diarrhea, dysentery and colitis (Kirtikar and Basu, 1975). The bark and leaves of the plant is reported to possess various medical uses such as anti-hepatotoxic (Kapil *et al*., 1995), antidiuretic, wound healing, antiseptic (Anonymous, 1992) and anthelminthic activity (Gunasekharan and Divyakant, 2006). Hence, the present study was formulated to find out the role of fruits of *N. cadamba* in reducing CaOx depositions in kidney.

## Materials and methods

### Collection and preparation of fruit extract

The fruits of *N. cadamba* were collected from the University Campus, Kariavattom (8°m37’36N, 76°50’14E), Thiruvananthapuram and was authenticated by the Department of Botany, University of Kerala, Kariavattom (Voucher No: KUBH 5811). The fruits were washed, dried, powdered and subjected to soxhlet extraction using methanol as solvent. The methanol fruit extract of *N. cadamba* (MFNC) was concentrated using rotary vacuum evaporator and the resultant extracts was stored in air tight containers.

### Experimental animal model

Healthy adult male albino rats of wistar strain weighing between 200- 250g were used for the anti-nephrolithiatic study. The animals were acclimatized to standard laboratory conditions and housed in polypropylene cages. They were maintained on 12-hr light and dark cycle and provided with regular standard rat feed (Sai Durga, Bangalore) and drinking water at *ad libitum*. The ethical clearance has been obtained from the institutional animal ethical committee prior to the experiment (Approval no: IAEC-KU-23/2011-12-ZOOL-GP (3)).

### Acute toxicity

The acute oral toxicity study was carried out as per the Organization for Economic Cooperation and Development (OECD) guidelines 423. Wistar albino rats were randomly selected and are grouped having three animals in each group. The fruit extract of *N. cadamba* was administered orally at one of the defined doses using a stomach tube. After the substance has been administered, feeding was suspended for further 3-4 hours. The doses viz. 5, 50, 300 and 2000 mg/kg body weight of AFNC and was observed for the 24hr mortality as suggested by Ecobichon (1997).

### Pharmacological screening for anti-nephrolithiatic activity

#### Induction of nephrolithiasis and study protocol

In the present study, calcium oxalate nephrolithiasis was induced in rats by free access to drinking water containing 0.75% ethylene glycol (EG) and 2% ammonium chloride (AC) for 28 days (Touhami *et al*., 2007). Animals were divided into nine groups containing four rats in each group. Group 1 served as normal control and received regular standard rat food and drinking water at *ad libitum*. EG and AC in drinking water was given to group II - IX for the induction of renal calculi till the 28^th^ day. Group II were treated with the standard anti-urolithiatic drug, cystone (750mg/kg body weight). Dose for the methanol fruit extract was selected as 200 and 400mg/kg body weight. Group III was considered as lithiatic control where as group IV and V received the MFNC at a dose of 400mg/kg and 200mg/kg body weight respectively after treatment with EG+AC and served as post treatment groups (PR). Group VI-IX was taken as co treatment groups (CR). Among the co treatment regimes, group VI and VIII received aqueous fruit extract of *N. cadamba* (400mg/kg and 200mg/kg body weight respectively) from 15^th^ day till 28^th^ day. Group VII and IX received MFNC (200mg/kg and 400mg/kg body weight respectively) from 1^st^ day till 28^th^ day. All drugs were given once daily by oral route using gastric tube.

### Assessment of lithiatic activity

#### Urine and serum analysis

During the experimental period, 24hr urine samples were collected on day 7, 14 and 28 were centrifuged and few drops of concentrated HCl was added to the supernatant and stored at 4°C. Supernatant was analyzed for protein (biuret method), phosphorus (UV‐ molybdate method), calcium (O-cresolphthalein-complexone method) and oxalate (Hodgkinson and Williams, 1972). Urinary sediment was strongly agitated and examined under high power (40x) objective of a microscope for the detection of abnormal constituents in urine like casts, crystals etc.

Blood was collected from all the experimental animals by heart puncture under anaesthetic condition. Serum was separated and analyzed for creatinine (alkaline picrate method), urea (diacetylmonoxime method), blood urea nitrogen (BUN) and uric acid (Caraway method).

#### Renal homogenate analysis and histopathological study

The isolated kidneys of the treatment groups were cleaned off to remove the extraneous tissues and one kidney from each animal was dried at 80°C in hot air oven. Dried kidneys were boiled in 1N HCl for 30 minutes and homogenized. The homogenate was centrifuged and analyzed for stone forming constituents such as phosphorous (Fiske and Subbarow, 1925), calcium (Lorentz, 1982), oxalate (Hodgkinson and Williams, 1972) and protein (Sadasivan and Manickam, 1996). Another kidney was preserved in 10% buffered formalin for histological observations. About 3-7 μm thickness sections of paraffin embedded kidney tissues were dewaxed in xylene, rehydrated in graded alcohol series and stained with haematoxylin at room temperature then counterstained with eosin. The haematoxylin eosin stained slides were observed under research microscope (LEICA) at 10x and 40x magnification and photomicrographs were taken. Pizzolato’s (1964) staining method was carried out for the detection of calcium oxalate crystals in kidney tissues using silver nitrate and hydrogen peroxide.

#### ICP-MS study of renal tissues

For sample preparation all the reagents used for the ICP-MS analysis were of suprapur grade (Merck, USA) and high purity water from Milli-Q water purification system (Thermo Scientific, Barnstead, Smart 2 pure) was used for dilution and preparation of sample. For the microwave digestion (Anton Paar, Multiwave 3000), 0.1g of the dried powdered kidney tissue was mixed with 69% HNO3 (7ml) and 30% H2O2 (1ml) and the vessel was immediately closed to avoid contamination. The samples were hold for 25min, zero ramp time at 350watts and then retained for 30 min, 5 ramp at 500 watts. The details of the instrument are Peristaltic pump speed - 40 rpm, Cool flow - 14.00 L/min, Spray time temperature - 2.7˚C, Sampling depth – 5, Plasma power - 1550 watts, Auxilary flow - 0.80 L/min, Nebulizer flow - 0.97 and No of main runs - 3.

#### Statistical analysis

The results were expressed as mean ± SEM. Statistical analysis was performed by one way analysis of variance (ANOVA) followed by Tukey multicomparison test and P values ≤ 0.05 were considered significant.

## Results

Acute toxicity study observed that all animals group could tolerate a maximum dose of 2000 mg/kg body weight without noticeable behavioural changes. Therefore, the LD50 cut off dose was found to be 2000mg/kg body weight for the fruit extract of *N. cadamba.* Hence, the 1/5 and 1/10 of the median lethal dose was fixed as effective doses as 400 mg/kg and 200 mg/kg body weight for the present study.

The chronic administration of 0.75% EG with 2% AC to male wistar rats revealed the increased urinary excretion of protein, phosphorous, oxalate and calcium in lithiatic control rats (Group III) when compared with the Group I (Table 1 and 2). All urinary parameters were increased significantly in calculi induced animals (lithiatic control) compared with control. The results clearly demonstrates a significant (p<0.01) decrease of protein in the curative regimes when compared to group III. Urinary protein concentration was found to have the same range when compared with the extract received groups and antilithiatic drug received groups. The *N. cadamba* fruit extract supplementation in post treatment regimes significantly reduced the levels of calcium. Urinary phosphorus concentration in control group is found to be 4.76±0.27 and is significantly (p<0.01) increased (15.15±1.68) in group III. When compared to the lithiatic control rats, phosphorus level in post treatment group was significantly low. A significant (p<0.01) decrease in urine phosphorus level was noticed from curative regimes when compared with group IV and V. The results of urinary parameters strongly suggest that the fruit extract can lower the risk factors of kidney stone *i.e.,* calcium, phosphorus and oxalate when compared to lithiatic rats.

**Table 1.**
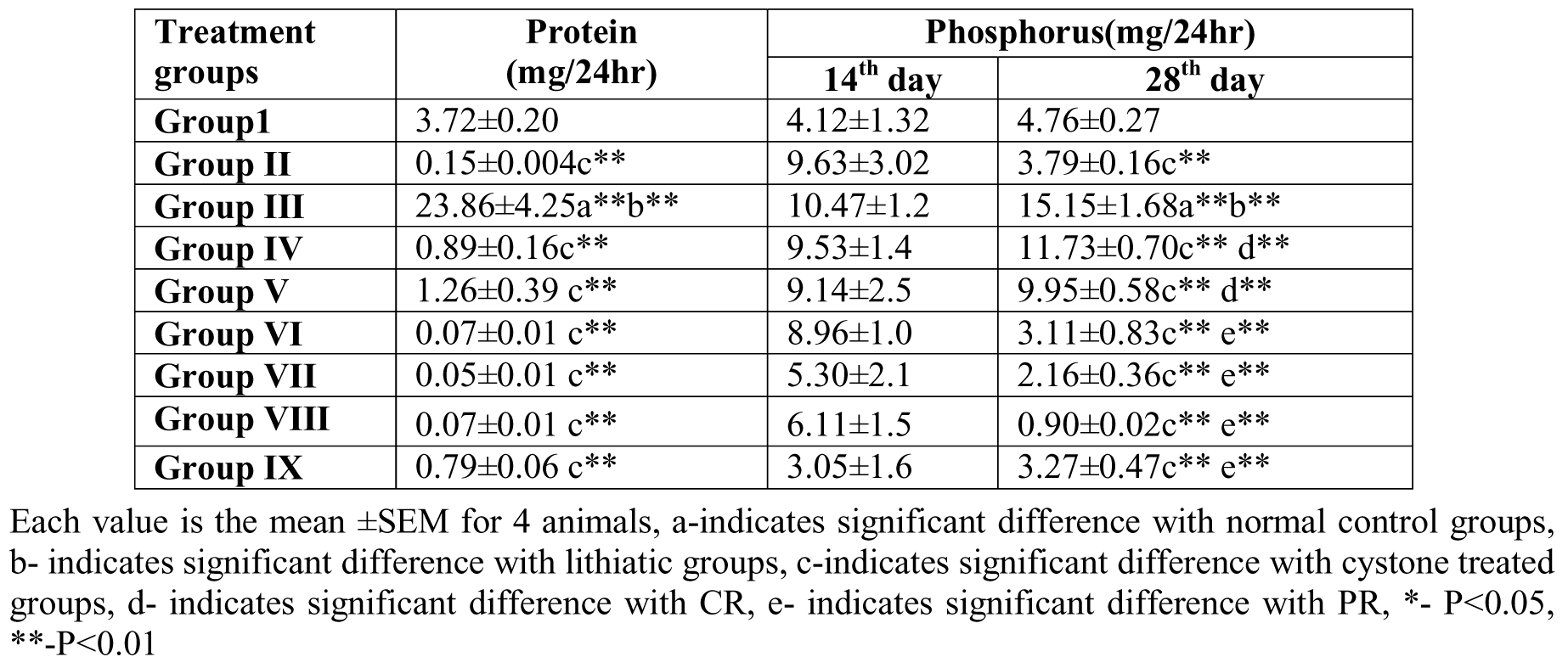
Effect of MFNC on urine protein and phosphorus

| Treatment groups | Protein (mg/24hr) | Phosphorus(mg/24hr) |  |
| --- | --- | --- | --- |
|  |  | 14<sup>th</sup> day | 28<sup>th</sup> day |
| **Group I** | 3.72±0.20 | 4.12±1.32 | 4.76±0.27 |
| **Group II** | 0.15±0.004c** | 9.63±3.02 | 3.79±0.16c** |
| **Group III** | 23.86±4.25a**b** | 10.47±1.2 | 15.15±1.68a**b** |
| **Group IV** | 0.89±0.16c** | 9.53±1.4 | 11.73±0.70c** d** |
| **Group V** | 1.26±0.39 c** | 9.14±2.5 | 9.95±0.58c** d** |
| **Group VI** | 0.07±0.01 c** | 8.96±1.0 | 3.11±0.83c** e** |
| **Group VII** | 0.05±0.01 c** | 5.30±2.1 | 2.16±0.36c** e** |
| **Group VIII** | 0.07±0.01 c** | 6.11±1.5 | 0.90±0.02c** e** |
| **Group IX** | 0.79±0.06 c** | 3.05±1.6 | 3.27±0.47c** e** |

Each value is the mean ±SEM for 4 animals, a-indicates significant difference with normal control groups, b- indicates significant difference with lithiatic groups, c-indicates significant difference with cystone treated groups, d- indicates significant difference with CR, e- indicates significant difference with PR, \*- P<0.05, \*\*-P<0.01

**Table 2.** Effect of MFNC on urine calcium and oxalate

| Treatment groups | Calcium (mg/24hr) |  | Oxalate (mg/24hr) |  |
| --- | --- | --- | --- | --- |
|  | 14<sup>th</sup> day | 28<sup>th</sup> day | 14<sup>th</sup> day | 28<sup>th</sup> day |
| **Group I** | 0.6±0.14 | 0.92±0.01 | 0.58±0.13 | 0.16±0.03 |
| **Group II** | 1.87±0.34 | 1.85±0.41 | 2.69±0.32 | 1.61±0.09 |
| **Group III** | 2.7±0.54 | 3.47±0.57 a* | 2.54±0.51 | 3.46±0.06a**b** |
| **Group IV** | 3.88±1.4 | 2.25±0.47 | 3.08±1.01 | 1.54±0.01c** |
| **Group V** | 3.46±0.7 | 1.45±0.04 | 3.45±1.10 | 1.13±0.01 c** |
| **Group VI** | 2.53±0.7 | 6.79±0.41 | 2.14±0.27 | 1.60±0.05 c** |
| **Group VII** | 4.51±1.7 | 8.52±0.7 | 1.9±0.41 | 2.20±0.35 c** |
| **Group VIII** | 3.83±1.3 | 6.59±0.27 | 2.50±0.31 | 1.01±0.003 c** |
| **Group IX** | 2.64± 1.5 e** | 2.03±0.54 | 1.66±1.05e** | 0.56±0.17 e** |

Each value is the mean ±SEM for 4 animals, a-indicates significant difference with normal control groups, b- indicates significant difference with lithiatic groups, c-indicates significant difference with cystone treated groups, d- indicates significant difference with CR, e- indicates significant difference with PR, \*- P<0.05, \*\*- P<0.01

Blood urea concentration has significantly (p <0.01) elevated in group III treated animals than control group of rats. However, these elevated levels are significantly decreased by the treatment with methanol fruit extract. Group VI (13.05±0.36) of co treatment has exhibited a remarkable decrease in nitrogenous wastes in blood. Both low and high doses of the MFNC seem to be effective in lowering the concentration of BUN in serum. From the results, it is evident that lithogenic rats elevated the stone forming constituent, uric acid significantly (P<0.01) when compared with normal control rats (9.58±1.08). The serum uric acid level has significantly declined in the post treatment groups of MFNC. When compared with the post treatment groups to co‐ treatment animals, group VI (2.86±0.27) of co treatment is found to have capacity to lower the uric acid concentration than any other groups. The results clearly demonstrate that high dose administered groups in both post and co treatments have significantly decreased the serum uric acid concentration. When compared to normal control rats (Group I), lithiatic control rats elevated the creatinine concentration in body fluids. But in co treated groups (Group VI to IX), the creatinine level was significantly (p<0.01) decreased, when compared with group V of post treatment regime. Significant (p<0.01) reduction in serum creatinine was noticed in group VII and the value is comparable to normal control, lithiatic control and antilithiatic control rats. The values are presented as table 3.

**Table 3.**
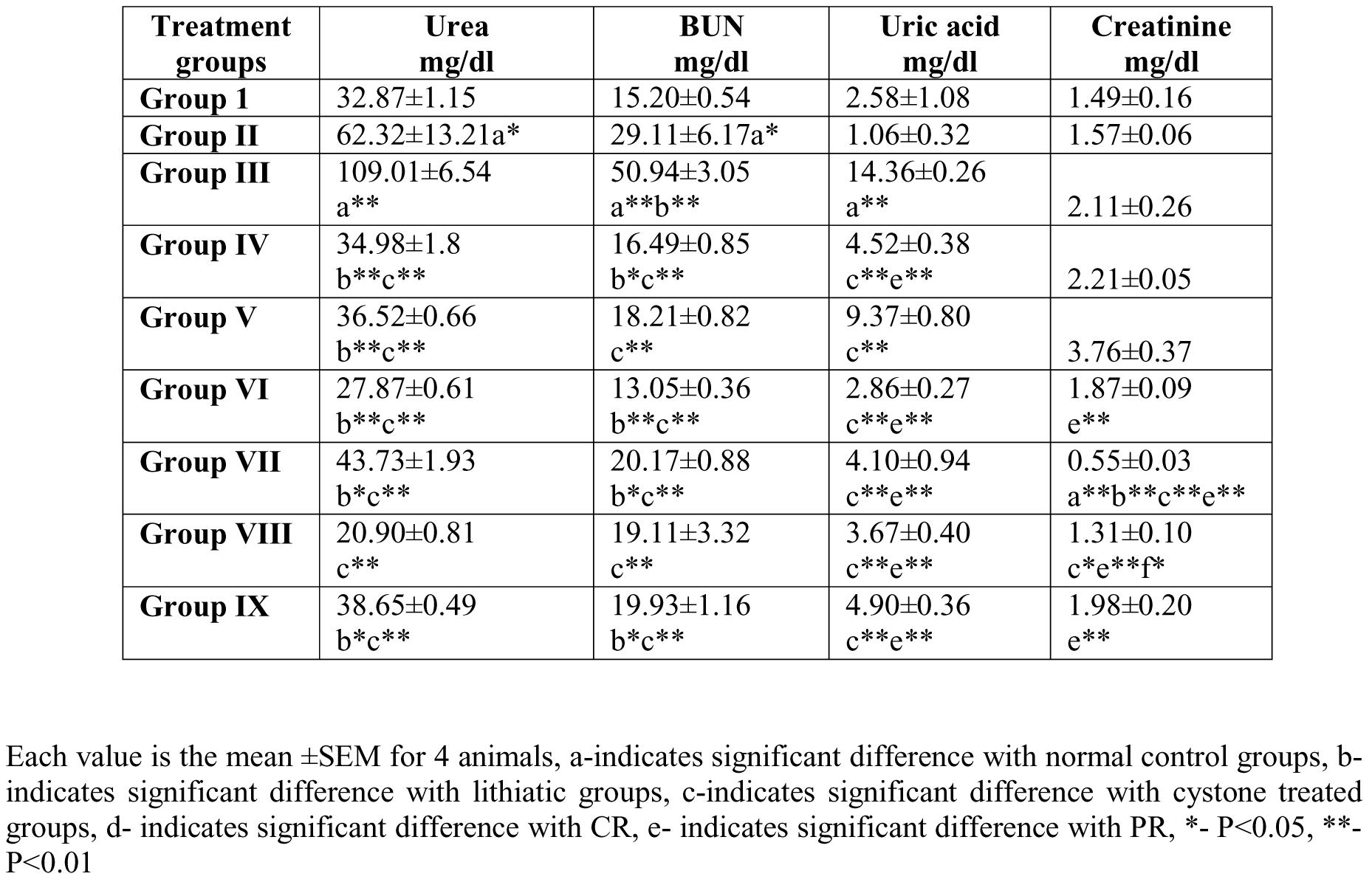
Effect of MFNC on serum biochemical parameters

Calcium, one of the main constituents of renal calculi is elevated significantly (p<0.01) in group III. The treatment with MFNC decreases the calcium deposition in kidneys significantly (p<0.01) than lithogenic rats. High dose supplemented animals in group IV (0.14±0.01) of post treatment has lowered the calcium concentration than low dose received groups of post treatment, group V (1.20±0.23). The present results have revealed the role of *N. cadamba* fruit extract in reducing the elevated kidney calcium concentration. Kidney oxalate values are found to be higher in urolithiatic rats (3.53±0.10) and is significant (p<0.01) when compared with group I. However, administration with MFNC has been significantly (p<0.01) reduced the level of oxalate in kidneys and the values are in the range of control animals. In both treatment regimes, group VII (0.37±0.14) and group IX (0.37±0.03), showed minimum oxalate concentration than any other groups. From the values, it is evident that both low dose (200mg/kg b. wt.) and high dose (400mg/kg b. wt.) of the fruit extracts are effective in reducing kidney oxalate concentration. The results have demonstrated that MFNC can significantly lower kidney phosphorus concentration than cystone treated animals. When compared with post treatment groups, the co treatment regimes are found to be effective in lowering phosphorus levels in kidneys (Table 4).

**Table 4.** Effect of MFNC on kidney

| Treatment groups | Protein (mg/24hr) | Phosphorus (mg/24hr) | Calcium (mg/24hr) | Oxalate (mg/24hr) |
| --- | --- | --- | --- | --- |
| Group I | 5.003±0.393 | 2.54±0.424 | 0.62±0.63 | 0.15±0.01 |
| Group II | 5.373±0.037 | 14.77±1.297a* | 1.06±0.21c** | 2.92±0.32a** |
| Group III | 4.733±0.296 | 3.82±0.109b** | 12.47±0.83 | 3.53±0.10a** |
| Group IV | 4.667±0.328 | 5.62±0.459 b** | 0.14±0.01a**c** | 0.73±0.28b**c** |
| Group V | 5.033±0.095 | 5.03±0.439 b** | 1.20±0.23 a**c** | 0.42±0.19 b**c** |
| Group VI | 5.090±0.231 | 3.86±0.204 b** | 0.89±0.17 a**c** | 0.70±0.13 b**c** |
| Group VII | 4.800±0.404 | 3.69±0.081b** | 0.27±0.24 a**c** | 0.37±0.14b**c** |
| Group VIII | 5.060±0.240 | 5.44±0.068 b** | 1.19±0.04 a**c** | 0.47±0.04 b**c** |
| Group IX | 4.900±0.252 | 4.82±0.313 b** | 1.84±0.38 a**c** | 0.37±0.05b**c** |

Each value is the mean ±SEM for 4 animals, a-indicates significant difference with normal control groups, b- indicates significant difference with lithiatic groups, c-indicates significant difference with cystone treated groups, d- indicates significant difference with CR, e- indicates significant difference with PR, \*- P<0.05, \*\*- P<0.01

Data obtained from the ICP-MS analysis exhibited a high kidney Mg concentration in group VI. Post treatment groups were observed with Mg value of 315.96 and 284.18. High Zn and Cu concentration was observed in CaOx stone forming control rats. Extract supplementation lowered its values in experimental rats. Zn concentration decreased more in group supplemented with MFNC at its low dose i.e. group VIII (31.09ppm). Group V (23.62ppm) was recorded with low level of Cu when compared with other treatment groups. When compared to group III, Fe concentration was lowered in MFNC treated groups. MFNC administration decreased the concentration of Al in experimental groups except in group VI, VII and IX. Group IV and V, the post treatment groups, and group VIII of co treatment groups showed decreased Al level. The test results were given in Table 5.

**Table 5.**
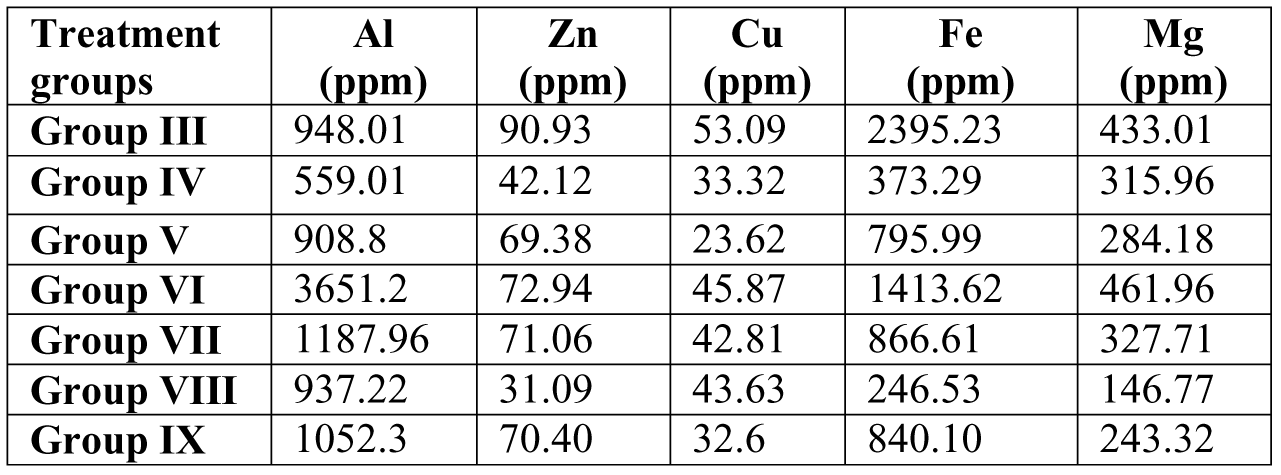
Effect of MFNC on elemental concentration in kidney

| Treatment groups | Al (ppm) | Zn (ppm) | Cu (ppm) | Fe (ppm) | Mg (ppm) |
| --- | --- | --- | --- | --- | --- |
| Group III | 948.01 | 90.93 | 53.09 | 2395.23 | 433.01 |
| Group IV | 559.01 | 42.12 | 33.32 | 373.29 | 315.96 |
| Group V | 908.8 | 69.38 | 23.62 | 795.99 | 284.18 |
| Group VI | 3651.2 | 72.94 | 45.87 | 1413.62 | 461.96 |
| Group VII | 1187.96 | 71.06 | 42.81 | 866.61 | 327.71 |
| Group VIII | 937.22 | 31.09 | 43.63 | 246.53 | 146.77 |
| Group IX | 1052.3 | 70.40 | 32.6 | 840.10 | 243.32 |

The urine microcrystal examination of all the treated groups of rats exhibited decreased crystalluria than ethylene glycol treated group. The photomicrograph of urine sample of post treated groups was observed with sparingly located dumb-bell shaped calcium oxalate monohydrate crystals, calcium oxalate dihydrate crystals and triple phosphate. But in group IV the number and size of the stones were decreased than in group V. The urine sample was observed with calcium oxalate monohydrate, dihydrate, struvite stones and uric acid stones. Group VI rats were reported with calcium oxalate monohydrate, dihydrate, uric acid and phosphate stones and the treatment with MFNC reduced the amount of stones. The day 7 urine sample of group VII observed with small sized stones whereas in the day 14, was noticed with less number of calcium oxalate stones by the administration of extract. MFNC reduced the crystal deposition in urine sample of day 28. Group VI is more effective in reducing the crystalluria than group VII received 200mg/kg b. wt. of MFNC. Group VIII exhibited small and aggregate stones of varying sizes. The observed stones from day 7 were calcium oxalate monohydrate, dihydrate and phosphate stones. After the supplementation with MFNC on day 28, the crystalluria were reduced in the urine samples. When compared to group VIII, group IX showed reduced amount of crystalluria. The calcium oxalate monohydrate, struvite and phosphate stones which are observed in day 14 were absent in urine sample at day 28.

When compared the histological sections of MFNC administered groups with lithiatic control rats, the fruit extract reduced the amount of stone depositions in all the treatment groups. Histopathological results clearly showed that the renal integrity and normal kidney architecture was found to be regained by MFNC in both post and co treatment groups. Special staining of the kidney tissues of all MFNC administered rats strengthens the observations in histopathology. In post treated groups, group V showed aggregated stone deposition in the region of renal tubules whereas in group IV, the stones were small in size and located in the glomerulus and in renal epithelial cells. In co treated groups, the amount of deposition of calcium oxalate stones was very less in group VI when compared to group VII in which stones were dispersed all regions of renal tissue. In group VIII the stones were concentrically arranged around the renal tubules and are seen as black dots and some stones were seen inside the tubules also. Group IX, received the highest dose of MFNC was noticed with less depositions of stones in the lumen as well as in the luminal epithelium of kidney when compared to all other co treated groups of rats.

**Fig.1.**
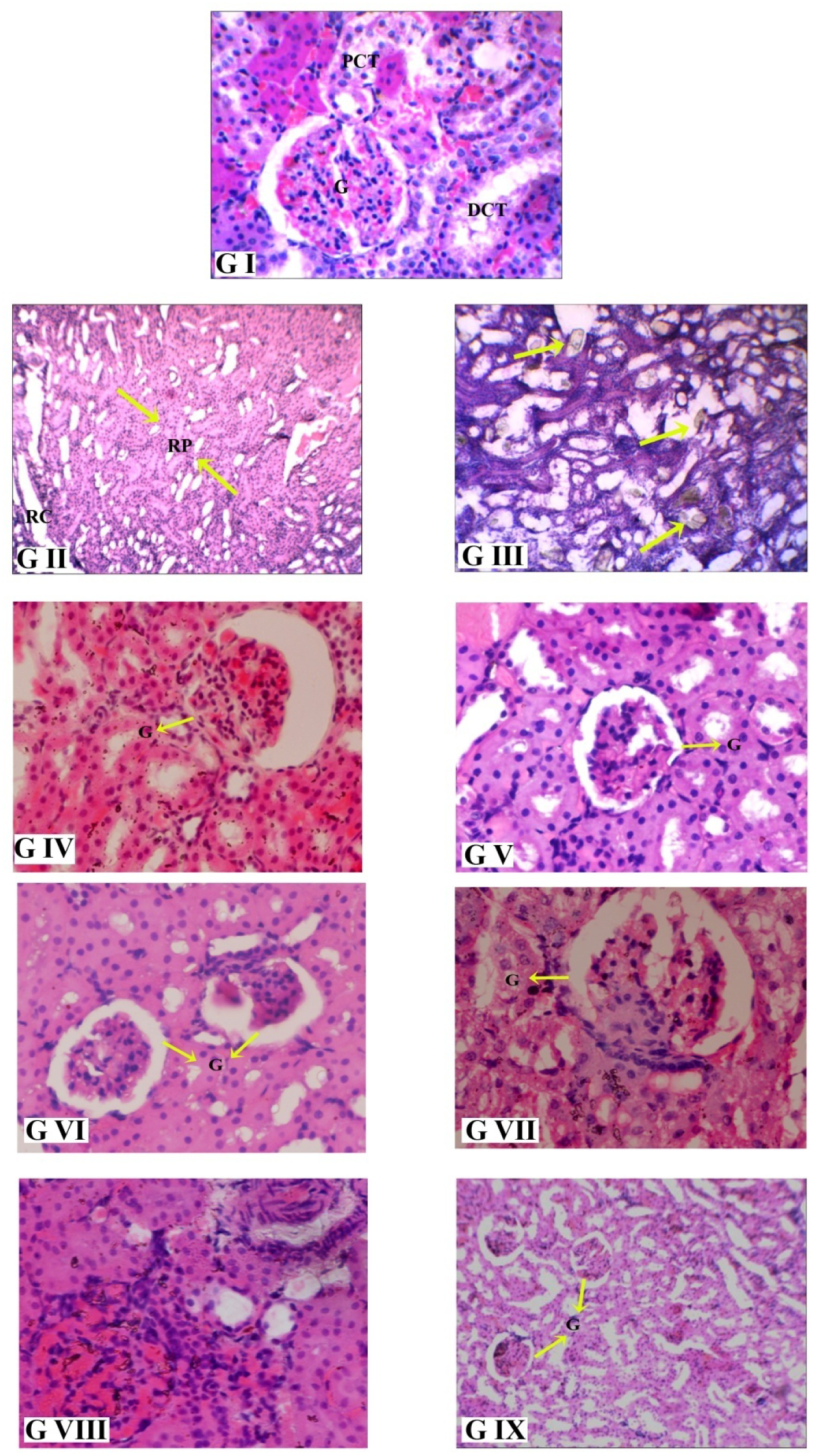
Histology of kidney tissues of experiments rats. G - Glomerules, PCT - Proximal Convoluted Tubuke, Dct - Distal Convoluted Tubule, RP‐ Rental Pyramid, RC - Rental Calyx

**Fig.2.**
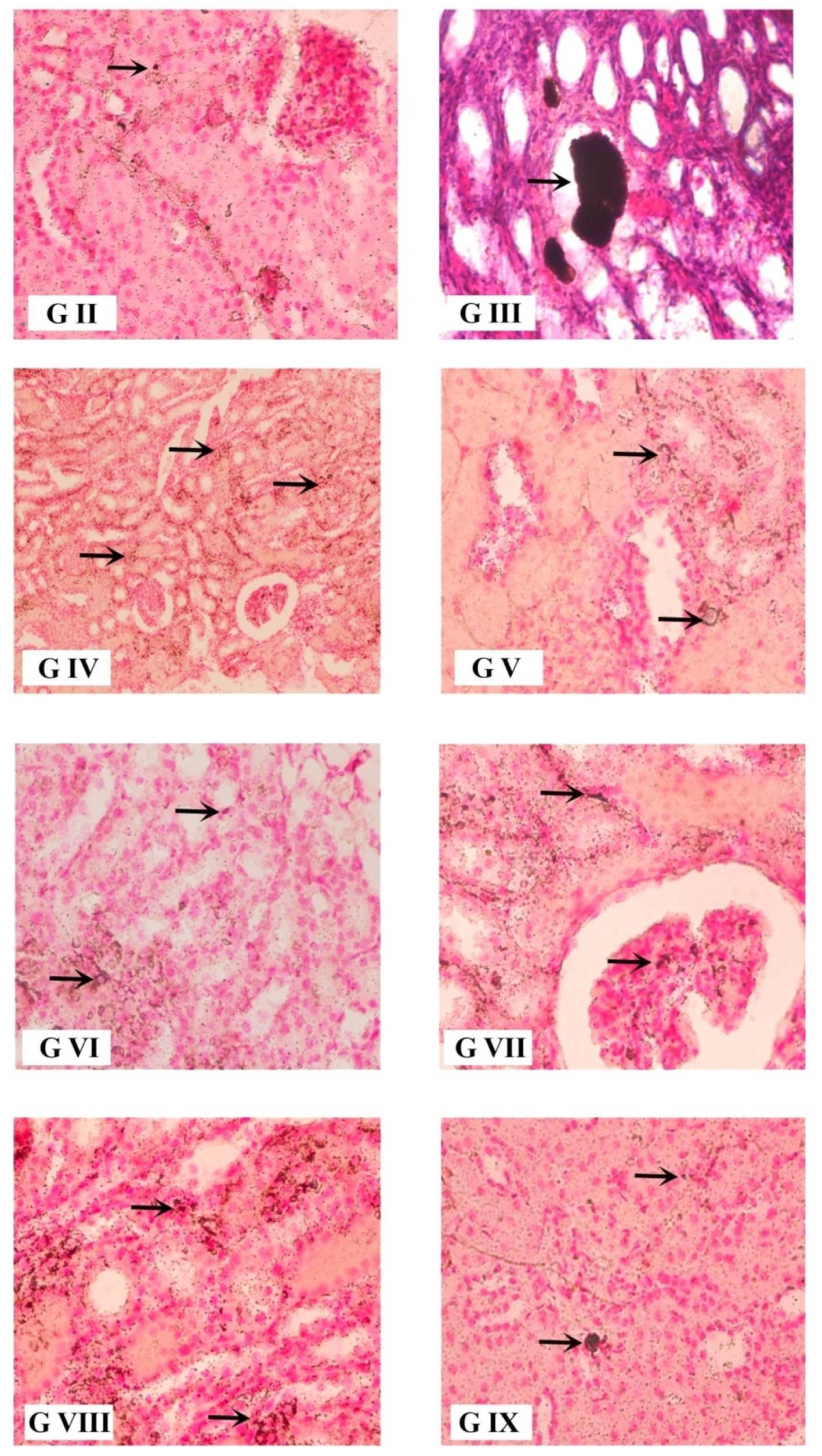
Pizzolato's staining of kidney tissues of experimental rats, → represents area of stone formations

**Fig.3.**
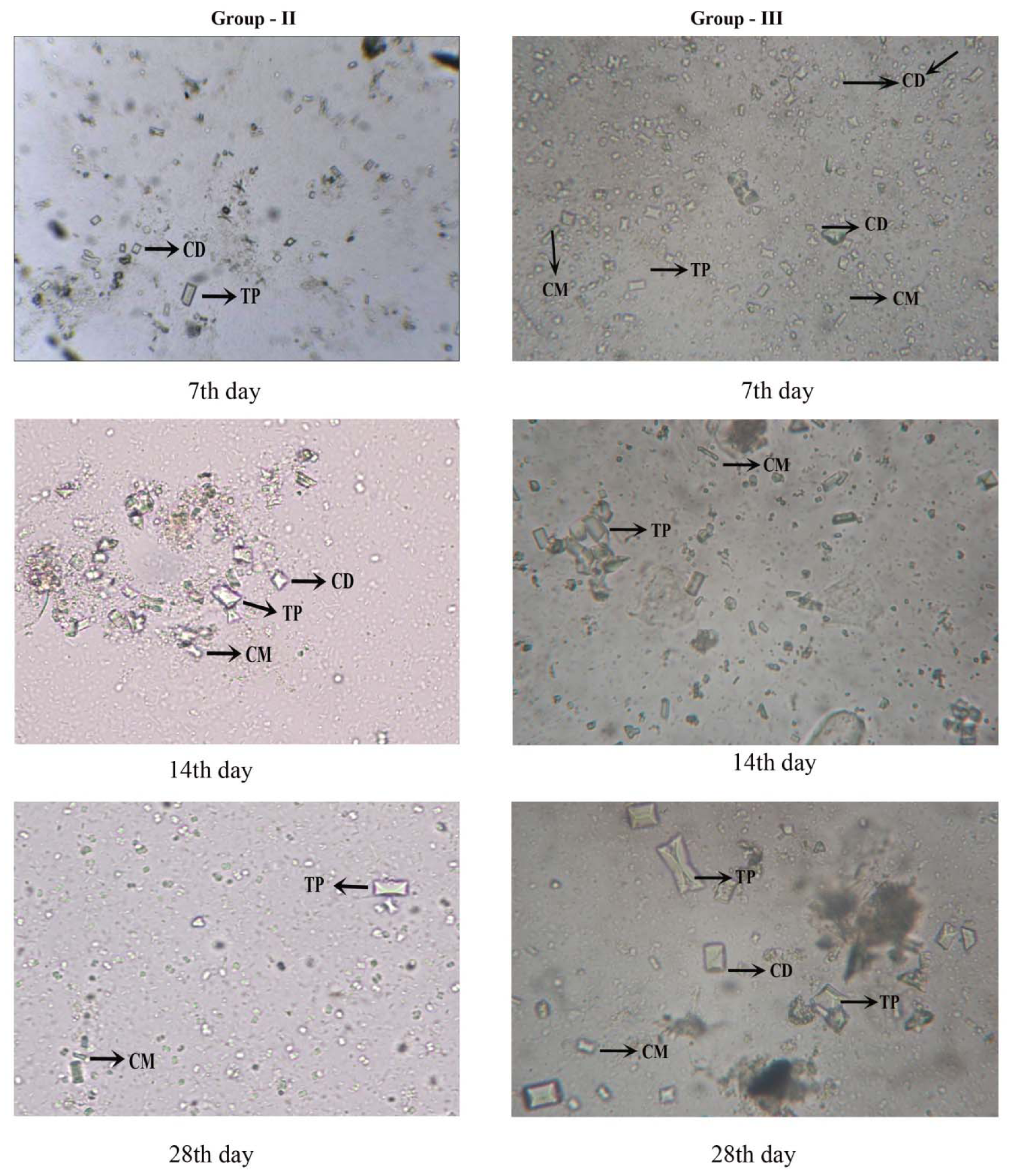
Urine micro crystal analysis of antilithiatic control and lithiatic control rats on day 7, 14 and 28, CD - Calcium oxalate Dihydrate, CM ‐Calcium oxalate Monohydrate TP‐ Triple Phosphate

**Fig.4.**
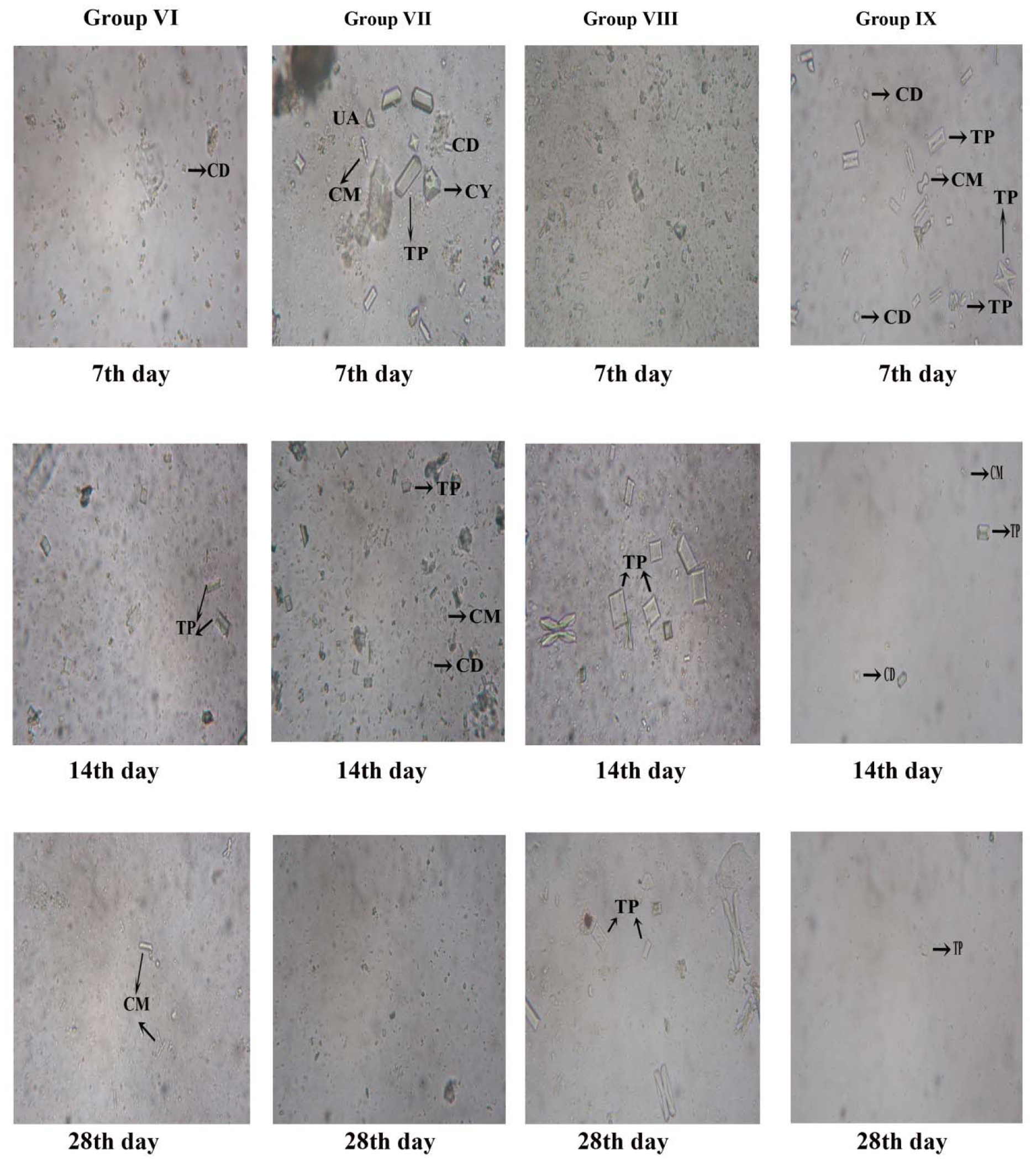
Urine micro crystal analysis of groups VI-IX experimental rats on day 7, 14 and 28, CD - Calcium oxalate Monohydrate, CD ‐Calcium oxalate Dihydrate TP‐ Triple Phosphate, CY - Cystein, UA - Uric Acid

## Discussion

EG /AC is generally used to mimic the renal calcium oxalate stone formation and deposition in humans (Atmani *et al*., 2003; Tsai *et al*., 2008). Here, fruit extracts of *N. cadamba* has been explored to evaluate its urinary protection in standard nephrolithiatic model. The stone inducing agents have increased the urinary oxalate and calcium in group III (lithiatic control) treated with EG/AC. The observed results are in agreement with previous studies which indicated that EG administration develops renal calculi which composed mainly of calcium oxalate (Selvam *et al*., 2001).

Hyperoxaluria is an important risk factor favouring renal calcium oxalate pathogenesis than hypercalciuria. The increased urinary calcium enhance the process of nucleation and precipitation of calcium and oxalate which causes calcium oxalate or calcium phosphate stones (Robert and Carman, 1963; Lehman *et al*., 1991). The lowered oxalate excretion is observed from the present study by the administration of fruit extract especially in group IX thus the extract has the capacity to lower the risk of hyperoxaluria. The urinary calcium has been reduced extensively in the group V and IX of the fruit extract supplemented ones. The findings are evident for the effectiveness of the extract in preventing calcium oxalate stone formation and aggregation. Here, the elevated phosphorus excretion in lithiatic control rats has brought back to its normal level through the supplementation of methanol fruit extract. The decreased calcium and phosphorus levels in methanol fruit extract of *N. cadamba* (MFNC) is evident for the action of the extract in curing calcium phosphate stones. Findings of previous studies have showed that the elevated oxalate and phosphate excretion induced the formation of calcium phosphate stones which also favours the development of calcium oxalate stones (Roger *et al*., 1997). The restoration of phosphate concentration in the fruit extract supplemented groups revealed the low risk of stone formation. The results confirm the preventing property of methanol fruit extract in calcium oxalate and calcium phosphate excretion.

The results pointed out that urinary protein levels are significantly lower in treatment groups than untreated groups. Previous studies described that the organic matrix have high protein content and these proteins have directive role in stone formation (Boyce, 1968; Atmani, 2001). Several studies demonstrated that inter I) family proteins and bikunin are the major proteins in the stone matrix and presence of these proteins modulate the process of crystal aggregation. The lowered urinary protein concentration by the methanol extract may reduce the formation of stone matrix or crystal aggregation. The data revealed that fruit extract could regulate the proteins involved in hyperoxaluria and crystal deposition. The increased calcium and phosphate excretion could be due to defective tubular reabsorption in kidneys as reported by Varalakshmi *et al*. (1990).

Previous studies have suggested that kidney possess a defence system against calcium oxalate crystallization and is mainly driven by the proteins of IαI family (Atmani, 2001). The protein lowering effect of fruit extract in all the treatment groups could be due to the effect of MFNC in enhancing defence system of kidney stone formation. Kidney deposition of calcium oxalate and phosphate concentration has elevated significantly (p<0.01) in rats which received EG/AC. The increased levels of kidney calcium oxalate and calcium oxalate crystals in nephron can produce damages in the epithelial lining of renal tubules, which causes the production of free radicals and heterogeneous nucleation and aggregation (Hadjzadeh *et al*., 2007). The increased concentration of the kidney calcium and phosphate in EG/AC administrated groups could be due to the production of oxidative stress from the free radicals. The results of the present study suggests that methanolic fruit extract reduces the calcium and oxalate in kidney which clearly revealed the free radical scavenging property as well as stone reducing effect of *N. cadamba* fruits and demonstrated the therapeutic effect of *N. cadamba* methanolic fruit extract on nephrolithiasis. All the treatment groups (group IV to IX) have exhibited a marked decrease in phosphate excretion than EG +AC administered group of rats. It can be speculated that methanol fruit extract of *N. cadamba* could reduce the calcium phosphate stones associated with calcium oxalate stones and the reduced phosphate concentration in turn will reduce the risk of stone formation. Treatment with methanol fruit extract of *N. cadamba* significantly (p<0.01) restores the phosphate level in kidney.

Serum biochemical analysis revealed that urea, uric acid, BUN and creatinine were accumulated in the blood of lithiatic control rats significantly (p<0.01). The increased nitrogenous waste materials in calculi induced rats (EG/AC administered groups) are due to the reduced glomerular filtration rate (GFR) which obstruct the urine flow by the deposition and block renal tubules with stones as reported by Ghodkar (1994). The present study also demonstrated that methanolic extract reduces the serum nitrogenous waste either by increasing glomerular filtration rate or by reducing the deposition of calcium oxalate stones. The elevated levels of serum urea, uric acid, BUN and creatinine indicated a remarkable damage in kidney and urinary system (Nayeem *et al*., 2010). However, treatment with methanolic fruit extract effectively reduces the renal damage and it could be the reason for decreased concentration of serum nitrogenous wastes.

The trace elements necessary for the metabolic process are stored and excreted by kidney which may result in the development of urinary stones (Keshavarzi *et al*., 2014). ICP-MS study revealed the presence of Al, Zn, Cu and Fe in high concentration in urolithiatic rats whereas; others (Welshman and Mc Geown, 1975., Sutor and Wooley, 1970) reported that Zn, Al, Co and Fe inhibited the precipitation of calcium oxalate. However, in the present study these elements were found to be increased in urolithiatic kidney. Trace elements (Zn, Cu, Ni and Fe) can produce weak soluble salts by combining with oxalate and phosphate ions and the same could be the reason for the development of kidney stones (Hesse and Siener, 2012). Infectious stones are incorporated by Zn phosphate as layers and the incorporation of trace elements into the urinary stone which depends on the type of stone and the occurrence of trace elements that affect the property of the stone (Hesse and Siener, 2012).

Magnesium (Mg) inhibits the process of super saturation of urine by preventing the absorption of oxalate. Decreased Mg concentration was observed in nephrolithiasis by increasing the level of excretion of oxalate. By this mechanism sufficient Mg is not available to form the Mg oxalate complex and thus reduce the incidence of stone formation (Deshmukh and Khan, 2006). This mechanism is attributed to the decreased kidney Mg level and increased urinary oxalate concentration in lithiatic control group (group III). The decreased kidney oxalate and increased urinary oxalate in MFNC administered groups could be the reason for the decreased Mg concentration in the treatment groups. Increased concentration of Fe content in group III may be the result of the trapping of Fe ions at the crystal surface or in the lattice as suggested by Hesse and Siener (2012). Inhibitory property of Cu on CaOx stone (Komleh *et al*., 1990) was not observed in the present study. The study suggests that Cu act as a promoter for crystal formation in kidney and elemental concentration significantly affected the mineralogical composition of stones. According to Munoz and Valiente (2005), precipitation of stone was increased with the concentration of Fe, Zn and Cu. The high concentration of Zn in the group III rat kidney is because of the increased precipitation of stone and MFNC administration decreased the level of Zn in kidney.

The histopathological studies demonstrated the presence of crystal depositions in renal cortex and renal medulla of lithiatic control rats. Tissue injury and inflammation in group III rat enhances the crystal retention, crystal adhesion and promotes stone formation (Thangarathinam *et al*., 2013). Supplementation of MFNC at 200mg/kg and 400mg/kg body weight revealed by the mild appearance of renal crystals which indicated the ability of the extract to dissolve the preformed renal stones and also prevent further stone formation (Laikangbam and Devi, 2012). MFNC administration improved the renal structure towards normal by reducing the oxalate and calcium concentration in kidney and reduces the super saturation of urine. These findings are in agreement with the reports of Christina and Muthumani (2013). Only few areas of calcification and small sized crystals and less renal tissue damage in the extract administered groups might be due to the action of the extract in the mineralization process of renal stone formation either in the growth or in the aggregation of the crystals (Saha and Verma, 2011). From all the above facts, it is evident that MFNC cured and prevented the growth of crystal depositions in treated groups (group IV-IX). Pizzolato’s staining exhibited positive results and stones were located in the lumen as well as in the epithelial lining of the tubules. Crystal depositions severely affected some areas and cause inflammation which leads to the loss of structural integrity and glomerular atrophied and the urinary space is occupied by crystals. The crystal deposition is due to the attachment of the crystals to the injured cells and are associated with the expression of crystal binding proteins like osteopontin (OPN), hyaluronic acid (HA) and CD44 (Yamate *et al*., 1998; Asselman *et al*., 2003).

## Conclusion

In the present study, EG/AC treatment through drinking water developed CaOx crystals in kidney and the administration of MFNC to urolithiatic rats reduced the deposition of crystals in urine and kidney and the extract is found to be effective to both curative and preventive treatment groups. Calcium oxalate crystals are found to be reduced by MFNC administration as evident from Pizzolato’s staining and histopathological study.

## Acknowledgement

The first author duly acknowledges the UGC-SAP facility of the Department of Zoology, University of Kerala, DST-PURSE Programme of University of Kerala and the financial assistance from UGC BSR.

